# Development and validation of IMMUNO-COV™: a high-throughput clinical assay for detecting antibodies that neutralize SARS-CoV-2

**DOI:** 10.1101/2020.05.26.117549

**Authors:** Rianna Vandergaast, Timothy Carey, Samantha Reiter, Patrycja Lech, Clement Gnanadurai, Mulu Tesfay, Jason Buehler, Lukkana Suksanpaisan, Shruthi Naik, Bethany Brunton, Jordan Recker, Michelle Haselton, Christopher Ziegler, Anne Roesler, John R. Mills, Elitza Theel, Scott C. Weaver, Grace Rafael, Matthew M. Roforth, Calvin Jerde, Sheryl Tran, Rosa Maria Diaz, Alice Bexon, Alina Baum, Christos A. Kyratsous, Kah Whye Peng, Stephen J. Russell

## Abstract

We here describe the development and validation of IMMUNO-COV™, a high-throughput clinical test to quantitatively measure SARS-CoV-2-neutralizing antibodies, the specific subset of anti-SARS-CoV-2 antibodies that block viral infection. The test measures the capacity of serum or purified antibodies to neutralize a recombinant Vesicular Stomatitis Virus (VSV) encoding the SARS-CoV-2 spike glycoprotein. This recombinant virus (VSV-SARS-CoV-2-S-Δ19CT) induces fusion in Vero cell monolayers, which is detected as luciferase signal using a dual split protein (DSP) reporter system. VSV-SARS-CoV-2-S-Δ19CT infection was blocked by monoclonal α-SARS-CoV-2-spike antibodies and by plasma or serum from SARS-CoV-2 convalescing individuals. The assay exhibited 100% specificity in validation tests, and across all tests zero false positives were detected. In blinded analyses of 230 serum samples, only two unexpected results were observed based on available clinical data. We observed a perfect correlation between results from our assay and 80 samples that were also assayed using a commercially available ELISA. To quantify the magnitude of the anti-viral response, we generated a calibration curve by adding stepped concentrations of α-SARS-CoV-2-spike monoclonal antibody to pooled SARS-CoV-2 seronegative serum. Using the calibration curve and a single optimal 1:100 serum test dilution, we reliably measured neutralizing antibody levels in each test sample. Virus neutralization units (VNUs) calculated from the assay correlated closely (p < 0.0001) with PRNT_EC50_ values determined by plaque reduction neutralization test against a clinical isolate of SARS-CoV-2. Taken together, these results demonstrate that the IMMUNO-COV™ assay accurately quantitates SARS-CoV-2 neutralizing antibodies in human sera and therefore is a potentially valuable addition to the currently available serological tests. The assay can provide vital information for comparing immune responses to the various SARS-CoV-2 vaccines that are currently in development, or for evaluating donor eligibility in convalescent plasma therapy studies.

## Introduction

At the end of 2019, a novel coronavirus, now named severe acute respiratory syndrome coronavirus 2 (SARS-CoV-2), emerged from Wuhan, China. The virus causes a variety of signs and symptoms, including most commonly cough, fever, and shortness of breath. While some individuals are asymptomatic or experience only mild illness, many experience severe symptoms and require hospitalization. SARS-CoV-2 spread quickly around the globe and was declared a global pandemic on March 11, 2020, by the World Health Organization. As of May 26, 2020, the virus has infected over 5.5 million people worldwide causing more than 346,000 deaths. PCR assays that detect active SARS-CoV-2 infection are playing an important role in tracking disease spread, while serological tests that detect antibodies against SARS-CoV-2 are being used to measure past rates of infection and identify individuals who may be immune to SARS-CoV-2. However, the availability of serological tests that can specifically measure SARS-CoV-2-neutralizing antibodies is severely limited.

Generation of virus-neutralizing antibodies is essential for blocking subsequent viral infections, and the presence of neutralizing antibodies correlates with protective immunity following vaccination (1). Yet, only a subset of virus-specific antibodies are neutralizing (2). It is not currently understood how total antibody levels to the virus or even to the Spike protein relate to neutralizing activity for SARS-CoV-2, and improved testing of SARS-CoV-2 neutralizing antibody responses is urgently needed. Early convalescent plasma therapy trials have shown promising results (3–5), but screening of plasma donors is inadequate if acceptance criteria are based on total anti-SARS2-CoV-2 antibody levels and not specifically on neutralizing antibody levels. Likewise, efficacy studies of new SARS-CoV-2 vaccines must consider not only total antibody responses, but neutralizing antibody responses.

Laboratory assays to detect SARS-CoV-2-neutralizing antibodies have been developed using clinical isolates of SARS-CoV-2 (6–8), pseudotyped lentiviral vectors (9,10), pseudotyped Vesicular stomatitis viruses (VSVs) (11), and, most recently, recombinant VSVs (12,13). We describe here IMMUNO-COV™, a high-throughput, scalable clinical assay for the detection of SARS-CoV-2-neutralizing antibodies using a recombinant VSV engineered to express the SARS-CoV-2 spike protein in place of the VSV-G glycoprotein. Several groups have generated VSVs encoding the SARS-CoV-2 spike (S) glycoprotein, which are being explored as potential vaccine candidates (World Health Organization website) or used in laboratory assays and research (12,13). The SARS-CoV-2 spike protein binds to angiotensin converting enzyme-2 (Ace-2) on the cell surface to initiate virus entry (14–16). As the primary surface-exposed viral protein, the coronavirus spike is a major target of the host immune system (11,17–19). Blocking the interaction of spike with Ace-2 prevents virus entry, making spike the primary target of neutralizing antibodies (9,13,15). Therefore, detection of antibodies capable of neutralizing VSV expressing spike glycoprotein is expected to directly reflect the level of SARS-CoV-2-neutralizing antibodies present. Importantly, using spike-expressing VSV in place of SARS-CoV-2 significantly increases assay safety and scalability.

Our assay exploits the fusion phenotype of spike-expressing VSV by using a dual split protein (DSP) luciferase reporter system to quantitate virus-induced cell fusion and thereby the level of virus neutralization (Figure 1). The DSP system uses a chimeric split green fluorescent protein (GFP) and split Renilla luciferase (EF446136.1) (20,21). Fusion between two cell lines expressing complementary pieces of the split reporter facilitates reassociation of fully functional GFP and Renilla luciferase. Thus, luciferase activity can be used to measure virus-induced cell fusion in a high-throughput 96-well plate format. Here, we describe the development of IMMUNO-COV™, including optimization of assay conditions and validation of the final clinical assay. An excellent correlation was observed between our assay results, clinical symptoms, and results from other serological tests. In particular, quantitation of SARS-CoV-2 neutralizing antibody levels correlated closely with plaque reduction neutralization EC50% reduction values from an assay using a clinical isolate of SARS-CoV-2 (22). Thus, IMMUNO-COV™ accurately quantitates SARS-CoV-2-neutralizing antibodies, making it a valuable tool for assessing plasma therapies and immune responses to new SARS-CoV-2 vaccines.

**Figure 1:**
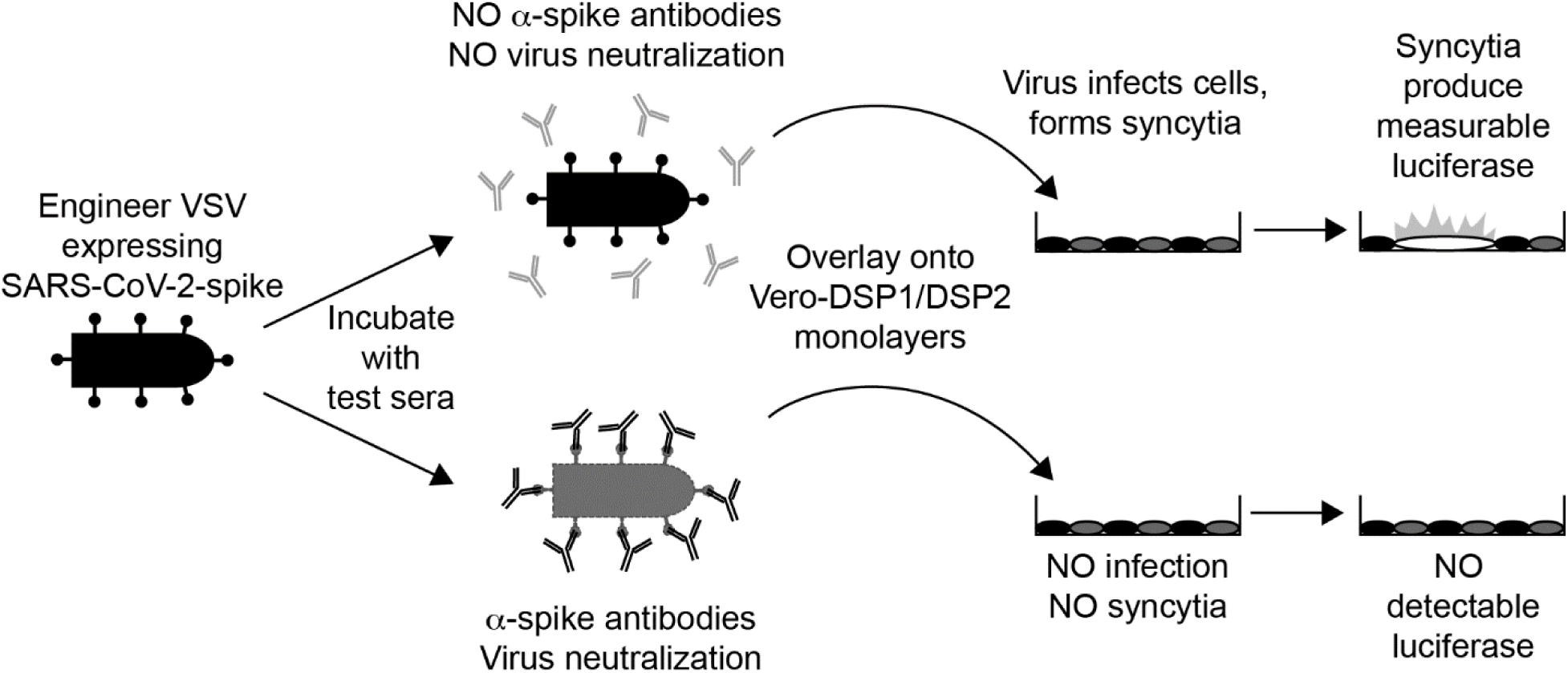
Overview of the IMMUNO-COV™ assay. A VSV expressing SARS-CoV-2 spike (VSV-SARS-CoV-2-S-Δ19CT) is incubated with test sera samples. In the absence of SARS-CoV-2-neutralizing antibodies (top) the virus retains infectivity and infects Vero-DSP1/DSP2 monolayers. If the test sample contains SARS-CoV-2-neutralizing antibodies (bottom), the antibodies bind to the spike protein causing virus neutralization by blocking cell entry. VSV-SARS-CoV-2-S-Δ19CT induces syncytia formation in Vero-DSP1/DSP2 monolayers, which reconstitutes a fully functional luciferase reporter that is used to quantitate virus-induced syncytia formation. High luciferase signal means the test sample did not neutralize the virus, while decreased luciferase indicates the presence of SARS-CoV-2-neutralizing antibodies in the test sample.

## Results

### Measurement of VSV-SARS-CoV-2-S-Δ19CT fusion through luciferase assay

Biosafety level 3 contaminant and practices are required to safely perform neutralization assays utilizing SARS-CoV-2, making widespread testing of patient samples using these assays impractical. We therefore sought to develop a safer, scalable assay that could be used to detect SARS-CoV-2 neutralizing antibodies in blood samples. To this end, we designed an assay in which syncytia formation in Vero cells, induced by infection with spike-expressing VSV, is detected using a dual split protein (DSP) luciferase reporter (Figure 1). Using reverse genetics, we generated a recombinant Vesicular stomatitis virus (VSV) in which the VSV glycoprotein (G) was replaced by the SARS-CoV-2 spike (S) protein (Figure 2A). Deletion of the last 19 amino acids in the C-terminal (cytoplasmic) domain of spike was required for efficient virus rescue and amplification. The resulting virus (VSV-SARS-CoV-2-S-Δ19CT), induced syncytia formation in Vero cell monolayers (Figure 2B). Virion incorporation of spike was confirmed in viral supernatants by immunoblot analysis (Figure 2C). Spike protein, but not VSV-G, was detected, confirming efficient replacement of VSV-G with SARS-CoV-2-S-Δ19CT in the recombinant virus.

**Figure 2:**
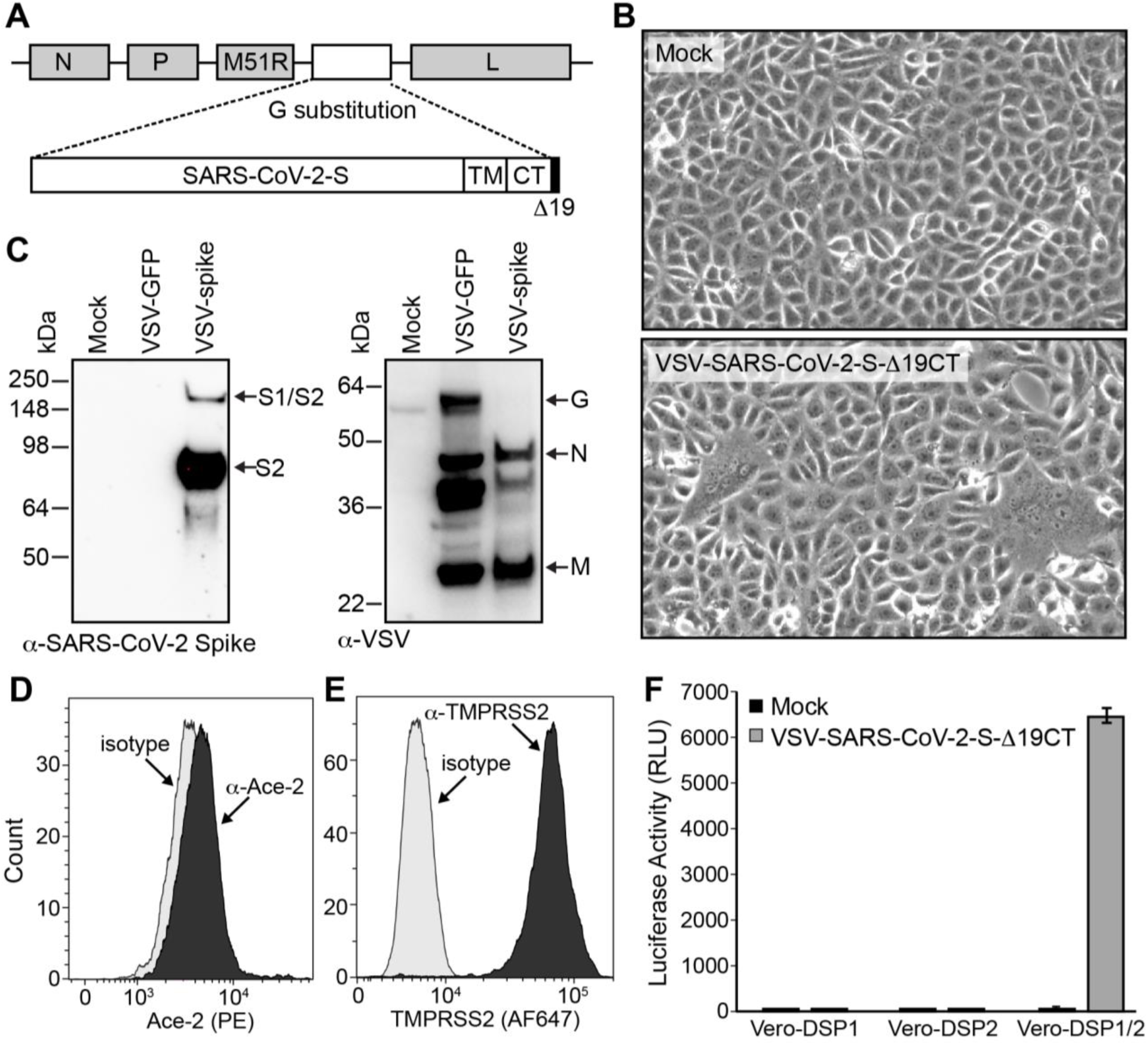
A luciferase assay to detect VSV-SARS-CoV-2-S-Δ19CT. A) Schematic representation of the VSV-SARS-CoV-2-S-Δ19CT genome. The location of the VSV N, P, M (M51R), and L genes are shown. In place of VSV-G a codon optimized SARS-CoV-2 spike gene with a 19 amino acid C-terminal (CT) deletion (Δ19CT) is substituted. TM is transmembrane domain. Not drawn to scale. B) VSV-SARS-CoV-2-S-Δ19CT induces syncytia formation. Vero monolayers were infected with VSV-SARS-CoV-2-S-Δ19CT or mock-infected. After 4 h, inoculums were removed and replaced with fresh media. Images were taken 20 h post infection at 100X magnification. C) Immunoblot analysis. Viral supernatants from mock-infected Vero cells or cells infected with VSV-GFP (encodes VSV-G) or VSV-SARS-CoV-2-S-Δ19CT were subjected to immunoblot analysis using α-SARS-CoV-2 spike antibody (left) and rabbit α-VSV antiserum (right). Arrows indicate the full-length S1/S2 variant and cleaved S2 variant of spike and the VSV-G, VSV-N, and VSV-M proteins. D and E) Flow cytometry. Monolayers of Vero-DSP1 cells were dislodged with versene, permeabilized (TMPRSS2 staining only), stained with isotype control (grey), α-Ace-2 (D, black) or α-TMPRSS2 (E, black) antibody, fixed, and subjected to flow cytometry analysis. F) VSV-SARS-CoV-2-S-Δ19CT induced luciferase activity. Monolayers of Vero-DSP1, Vero-DSP2, or mixed Vero-DSP/Vero-DSP2 cells were mock-infected or infected (10,000 TCID50 units/well) with VSV-SARS-CoV-2-S-Δ19CT. Luciferase activity was measured as a marker of cell fusion at 24 h after infection. Values represent the average (mean) RLU ± standard deviation from duplicate wells.

Since Vero cells were infected by VSV-SARS-CoV-2-S-Δ19CT and expressed both Ace-2 and TMPRSS2 (Figure 2D&E), they were chosen as the reporter cell line for our assay. Parental Vero cells were transduced with self-inactivating lentiviral vectors encoding DSP1 (DSP_1-7_) or DSP2 (DSP_8-11_) and a puromycin resistance gene, and reporter-expressing cells were selected using puromycin. When mixed monolayers of Vero-DSP1 and Vero-DSP2 cells seeded at a 1:1 ratio were infected with VSV-SARS-CoV-2-S-Δ19CT, we observed syncytia formation and readily detected luciferase activity at 24 h post infection (Figure 2F). In contrast, luciferase activity was not detected in infected monolayers of Vero-DSP1 alone or Vero-DSP2 alone, or in mock-infected Vero-DSP1/Vero-DSP2 monolayers. Thus, luciferase activity was specific to virus-induced cell fusion of mixed Vero-DSP1/Vero-DSP2 monolayers.

### Optimization of VSV-SARS-CoV-2-S-Δ19CT fusion

Syncytia formation is required to produce functional luciferase in our assay. However, complete destruction of the cell monolayer following syncytia formation reduces luciferase activity due to loss of cell viability. We therefore determined the optimal time after VSV-SARS-CoV-2-S-Δ19CT infection to measure luciferase. EnduRen™ Live Cell Substrate (EnduRen™) is an engineered “pro-substrate”, which must be cleaved by cellular esterases to produce the Renilla luciferase substrate coelenterazine. Because the pro-substrate is highly stable, luciferase activity can be measured repeatedly at any time between 2 and 24 hours after EnduRen™ addition. By 15 h after infection, luciferase activity in infected Vero-DSP1/Vero-DSP2 monolayers was readily distinguished from background signal in mock-infected cells (Figure 3A). Luciferase activity continued to rise, reaching peak levels around 26 hours after infection. Thus, optimal luciferase readout in the assay was between 24 and 30 hours after infection.

**Figure 3:**
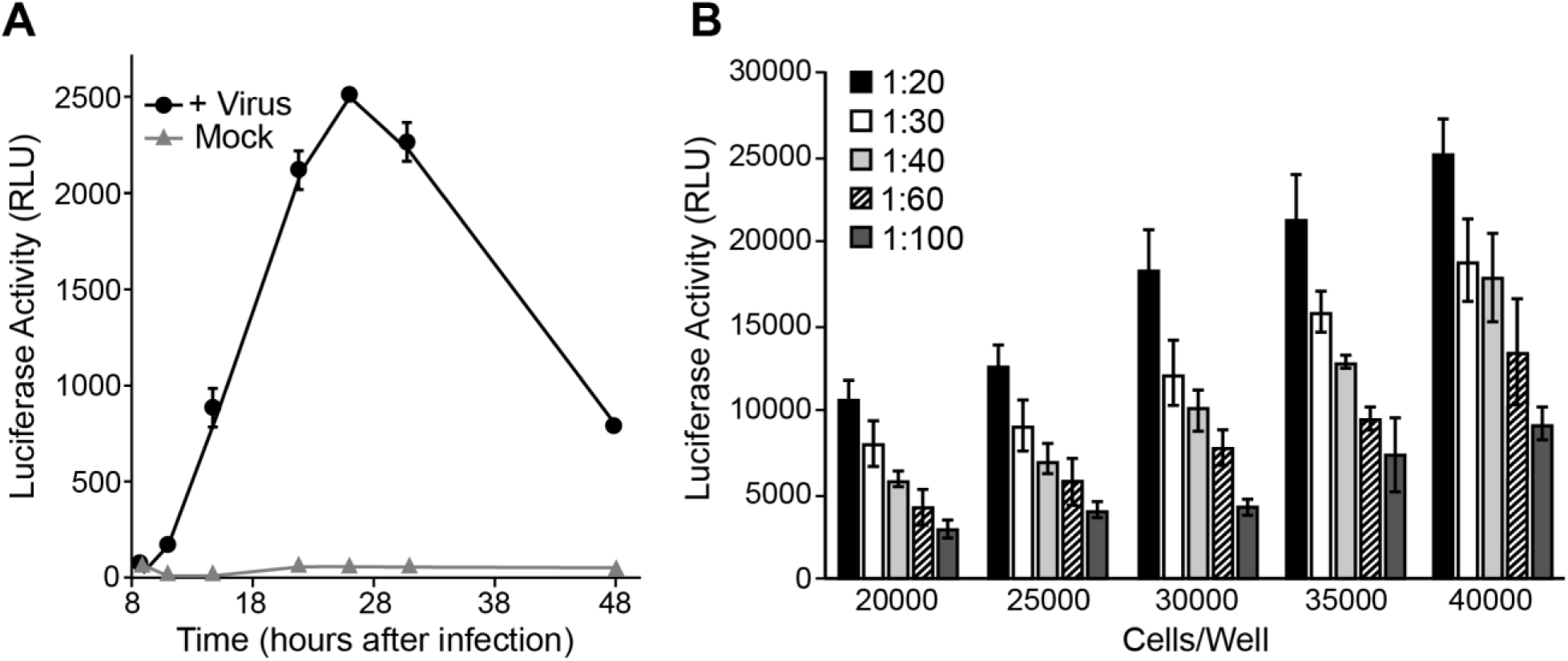
Optimization of luciferase signal. A) Kinetics of virus-induced luciferase activity. VSV-SARS-CoV-2-S-Δ19CT (2360 TCID50 units/well) or OptiMEM alone (mock) was overlaid onto monolayers of Vero-DSP1/Vero-DSP2 cells. Luciferase substrate EnduRen™ was added to wells and luciferase activity was measured at the indicated times (h) thereafter. Values represent the average (mean) RLU ± standard deviation from duplicate wells. B) Optimization of cell density. The indicated numbers of Vero-DSP1/Vero-DSP2 cells were seeded in 96-well plates. The following day, VSV-SARS-CoV-2-S-Δ19CT was diluted in OptiMEM to the indicated dilutions and overlaid onto the cell monolayers. EnduRen™ was added to wells and luciferase activity was measured 28 h after infection. Values represent the average (mean) RLU ± standard deviation from two experiments run in duplicate.

To enhance cell fusion and luciferase activity, and thereby increase assay sensitivity, we also determined the optimal cell density and virus concentration to use for assay. To this end, we seeded Vero-DSP1/Vero-DSP2 cells at increasing cell densities and after 24 hours infected the monolayers with increasing concentrations of VSV-SARS-CoV-2-S-Δ19CT. At all virus dilutions, increasing cell density improved luciferase signal (Figure 3B). We concluded that 4×10^4^ cells/well should be used for the assay. As expected, syncytia formation and luciferase activity also increased when more virus was used to inoculate the monolayers. To ensure lot-to-lot consistency between different virus preparations, an appropriate concentration of virus to use for assay was determined for each lot of virus based on performance against a set of contrived controls (see below).

### Detection of VSV-SARS-CoV-2-S-Δ19CT neutralization

To demonstrate that virus neutralization could be detected using our assay, we first tested the capacity of several purified proteins to inhibit virus-induced fusion. Affinity-purified polyclonal α-SARS-CoV-spike antibody and soluble recombinant Ace-2 protein bind spike on the virus surface and block entry into cells. Similarly, monoclonal α-Ace-2 antibody blocks the cellular receptors and limits virus entry. All three inhibitors successfully blocked VSV-SARS-CoV-2-S-Δ19CT-induced fusion in a dose-dependent manner (Figure 4A), demonstrating that our assay could detect inhibition of VSV-SARS-CoV-2-S-Δ19CT infection. Importantly, we also detected VSV-SARS-CoV-2-S-Δ19CT neutralization by plasma acquired from a SARS-CoV-2 convalescing individual known to have contracted and exhibited COVID-19 disease (referred to as NL1). Compared to pooled plasma from three presumed SARS-CoV-2 seronegative individuals, heat-inactivated NL1 plasma neutralized VSV-SARS-CoV-2-S-Δ19CT in a dose-dependent manner (Figure 4B), providing proof-of-concept for our assay.

**Figure 4:**
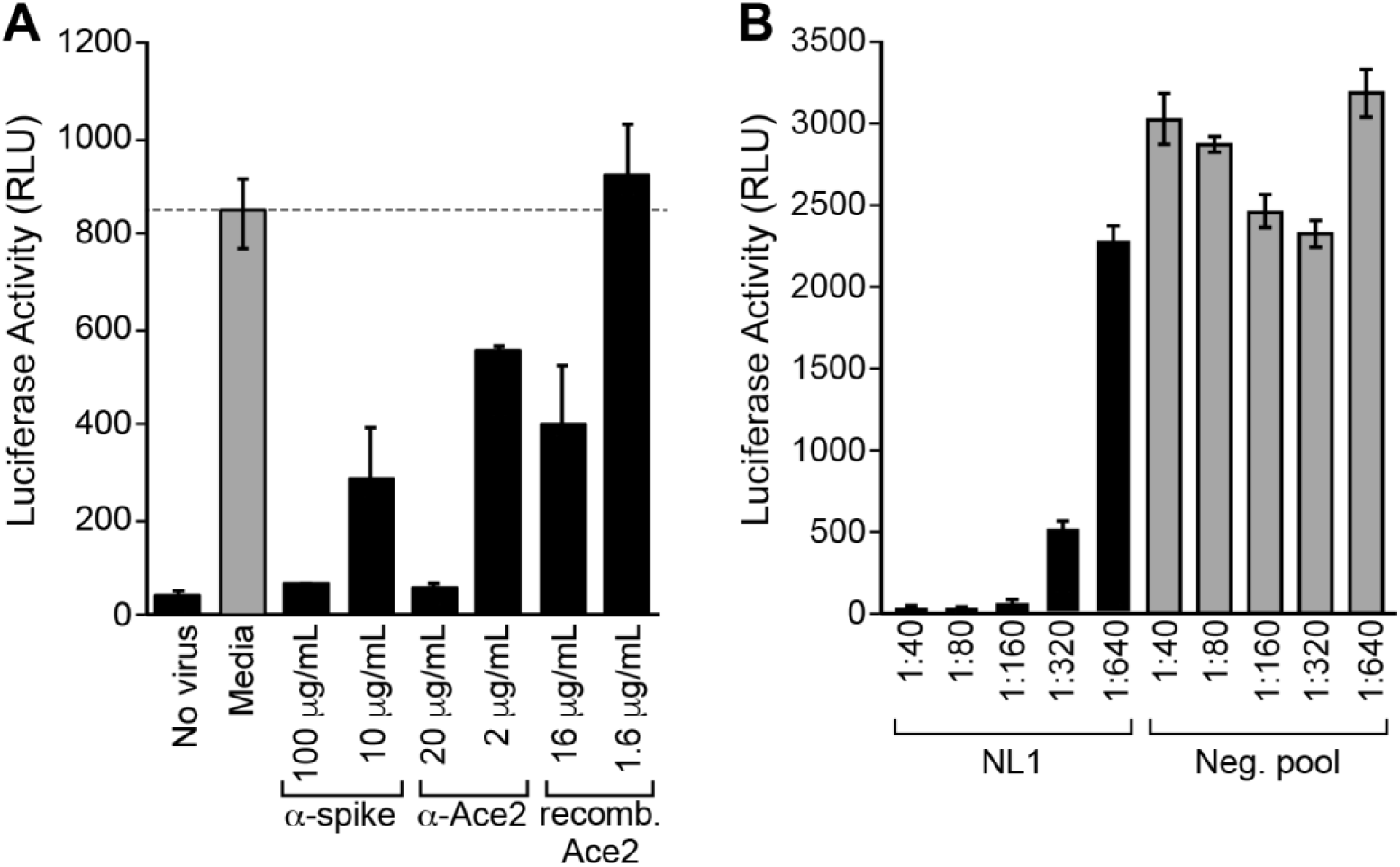
Neutralization of VSV-SARS-CoV-2-S-Δ19CT-induced luciferase activity. A) Inhibition by purified molecules. VSV-SARS-CoV-2-S-Δ19CT was incubated with media alone or the indicated concentrations of α-SARS-CoV-2-spike antibody, α-Ace-2 antibody, or recombinant Ace-2. After 1 h at 37°C, the virus mixes were overlaid on Vero-DSP1/Vero-DSP2 monolayers. Control wells received media alone (no virus). Luciferase substrate EnduRen™ was added to wells and luciferase activity was measured 24 h after infection. Values represent the average (mean) RLU ± standard deviation from duplicate wells. B) Inhibition by COVID-19 convalescing plasma. VSV-SARS-CoV-2-S-Δ19CT was incubated with the indicated dilutions of plasma from a COVID-19 convalescing individual (NL1) or pooled plasma from three presumptive SARS-CoV-2 seronegative individuals. Plasma dilutions represent final dilutions after addition of virus. After 1 h at 37°C, the virus mixes were overlaid on Vero-DSP1/Vero-DSP2 monolayers. EnduRen™ was added to wells and luciferase activity was measured 28 h after infection. Values represent the average (mean) RLU ± standard deviation from duplicate wells.

### Optimization of VSV-SARS-CoV-2-S-Δ19CT neutralization time

In an early experiment we observed that even in the absence of any neutralizing serum or inhibitors, incubation of VSV-SARS-CoV-2-S-Δ19CT at 37°C for 1 hour resulted in loss of luciferase activity (Figure 5A). This loss was partially abrogated at higher concentrations of virus. Nevertheless, we explored whether a shorter incubation time at room temperature was sufficient for virus neutralization. Two-fold serial dilutions of neutralizing NL1 plasma were prepared and incubated with VSV-SARS-CoV-2-S-Δ19CT for 60 minutes at 37°C or for 15, 30, or 60 minutes at room temperature. Relative to the other conditions, luciferase activity was slightly higher at the 1:160 dilution in the sample that had been incubated for 15 minutes at room temperature (Figure 5B), suggesting that this condition may not be sufficient for complete neutralization. However, NL1 neutralized VSV-SARS-CoV-2-S-Δ19CT as well, if not more efficiently, when incubated at room temperature for 30 or 60 minutes relative to incubation at 37°C for 1 hour. We concluded that a 30-minute incubation at room temperature is sufficient for virus neutralization.

**Figure 5:**
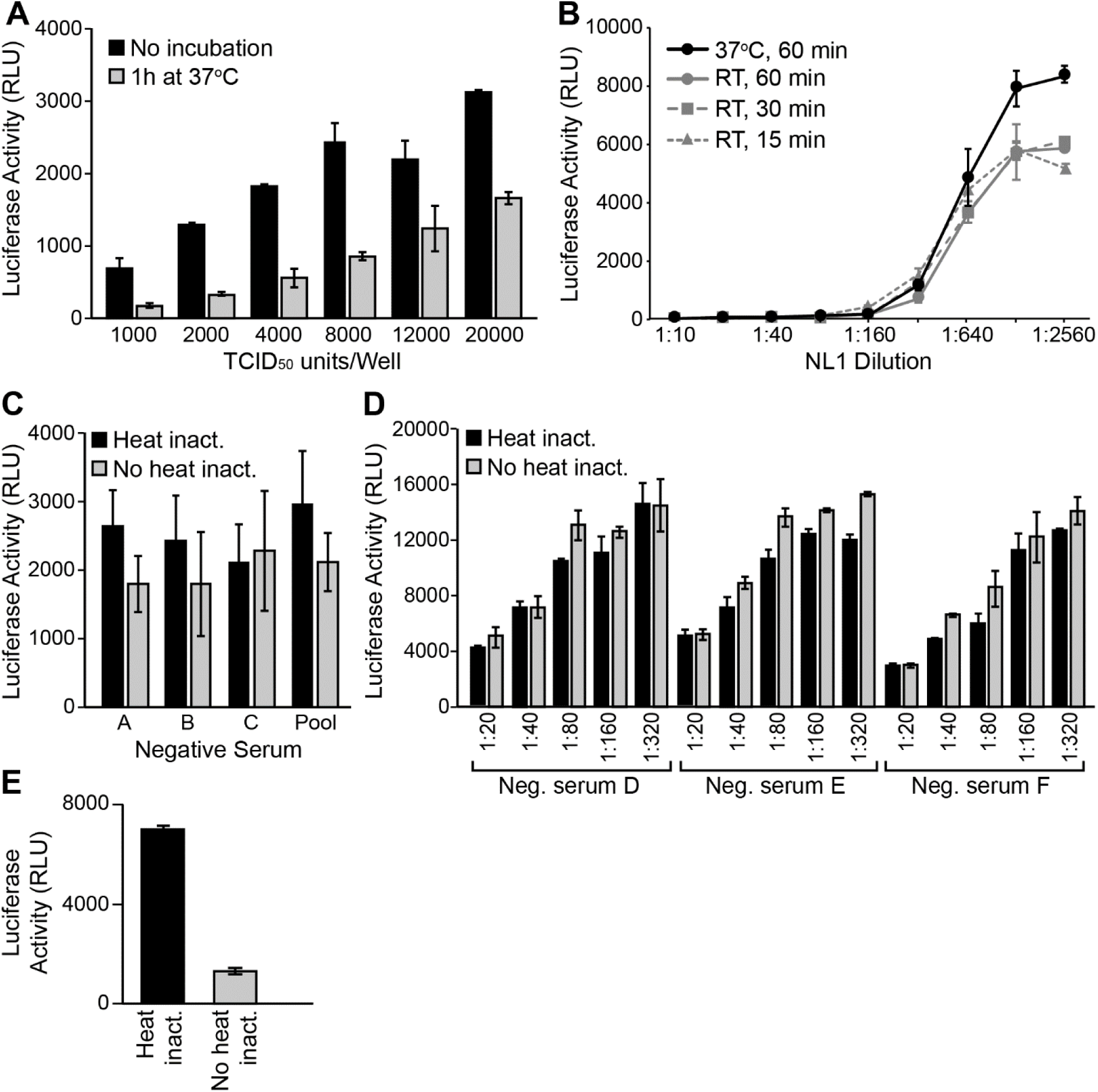
Optimization of neutralization conditions. A) Incubation of virus decreases fusion. VSV-SARS-CoV-2-S-Δ19CT was thawed and diluted to the indicated TCID_50_ units/100 µL in OptiMEM. Duplicate samples were prepared; one set was immediately overlaid onto Vero-DSP1/Vero-DSP2 monolayers, while the other was incubated at 37°C for 1 h before being overlaid onto Vero-DSP1/Vero-DSP2 monolayers. Luciferase substrate EnduRen™ was added to wells and luciferase activity was measured 27 h after infection. Values represent the average (mean) RLU ± standard deviation from duplicate wells. B) NL1 inhibition under different conditions. Dilutions of NL1 convalescing plasma were prepared and incubated with VSV-SARS-CoV-2-S-Δ19CT for the times and at the temperatures indicated before being overlaid onto Vero-DSP1/Vero-DSP2 monolayers. Dilutions represent NL1 dilutions in the final virus/plasma mixes. EnduRen™ was added to wells and luciferase activity was measured 25 h after infection. Values represent the average (mean) RLU ± standard deviation from duplicate wells. C&D) Effect of heat inactivation on negative sera. Presumptive negative serum samples A, B, C, D, E, F, and a commercial SARS-CoV-2 seronegative pooled sera (Pool) were prepared as duplicate aliquots. One aliquot was stored on ice while the other was incubated for 30 min at 56°C. Each aliquot was then diluted in OptiMEM and mixed with VSV-SARS-CoV-2-S-Δ19CT to a final dilution of 1:100 (C) or the dilutions indicated (D). Following a 30 min incubation at room temperature, the virus/sera mixes were overlaid onto Vero-DSP1/Vero-DSP2 monolayers. EnduRen™ was added to wells and luciferase activity was measured 23 h after infection. Values represent the average (mean) RLU ± standard deviation from duplicate wells. E) Effect of heat inactivation on COIVD-19 convalescing serum. A presumptive SARS-CoV-2 seropositive sample was thawed and assayed as described in Panel D at a 1:40 final dilution.

### Heat inactivation is not necessary for assay compatibility

To further streamline our assay workflow and reduce the burden of pre-analytical sample processing, we determined whether heat inactivation of samples is necessary to achieve reliable results. Serum samples were used for testing due to their greater availability compared to plasma samples. We initially tested three presumptive SARS-CoV-2 seronegative samples and a commercially available sera pool (generated before fall of 2019) at a 1:100 dilution. In the sera pool, and two of the three negative sera, heat-inactivation led to a modest decrease in luciferase activity (Figure 5C), possibly due to low-levels of virus inactivation by complement. Yet overall, heat-inactivation had little effect on VSV-SARS-CoV-2-S-Δ19CT-induced cell fusion. When three additional presumptive seronegative sera were tested at multiple dilutions, we again saw little difference in luciferase activity between heat-inactivated and non-heat-inactivated samples (Figure 5D). In contrast, we noted that in one sample heat inactivation reduced the capacity of a SARS-CoV-2 convalescing serum to neutralize VSV-SARS-CoV-2-S-Δ19CT (Figure 5E). We therefore concluded that using non-heat inactivated samples is preferred, as it streamlines assay workflow and increases assay sensitivity.

### Determination of the minimum recommended dilution

Sample interference is common in serological assays at high sample concentrations. Therefore, we sought to define the minimum recommended dilution for assay samples, in which false positives due to sample interference are nearly eliminated. To this end, we assayed thirty-nine presumptive SARS-CoV-2 seronegative samples at 2-fold serial dilutions ranging from 1:10 through 1:320. As expected, interference with virus fusion and luciferase activity was observed at higher serum concentrations (Figure 6A), but disappeared as the samples were further diluted. Statistical analyses indicated that the minimum recommended dilution was approximately 1:100. At this dilution there is greater than 95% confidence that the luciferase signal from a negative serum sample will be ≥ 50% relative to a pooled SARS-CoV-2 seronegative sera control. In all future studies, therefore, we used 1:100 as the dilution for both test serum samples as well as pooled SARS-CoV-2 seronegative controls.

**Figure 6:**
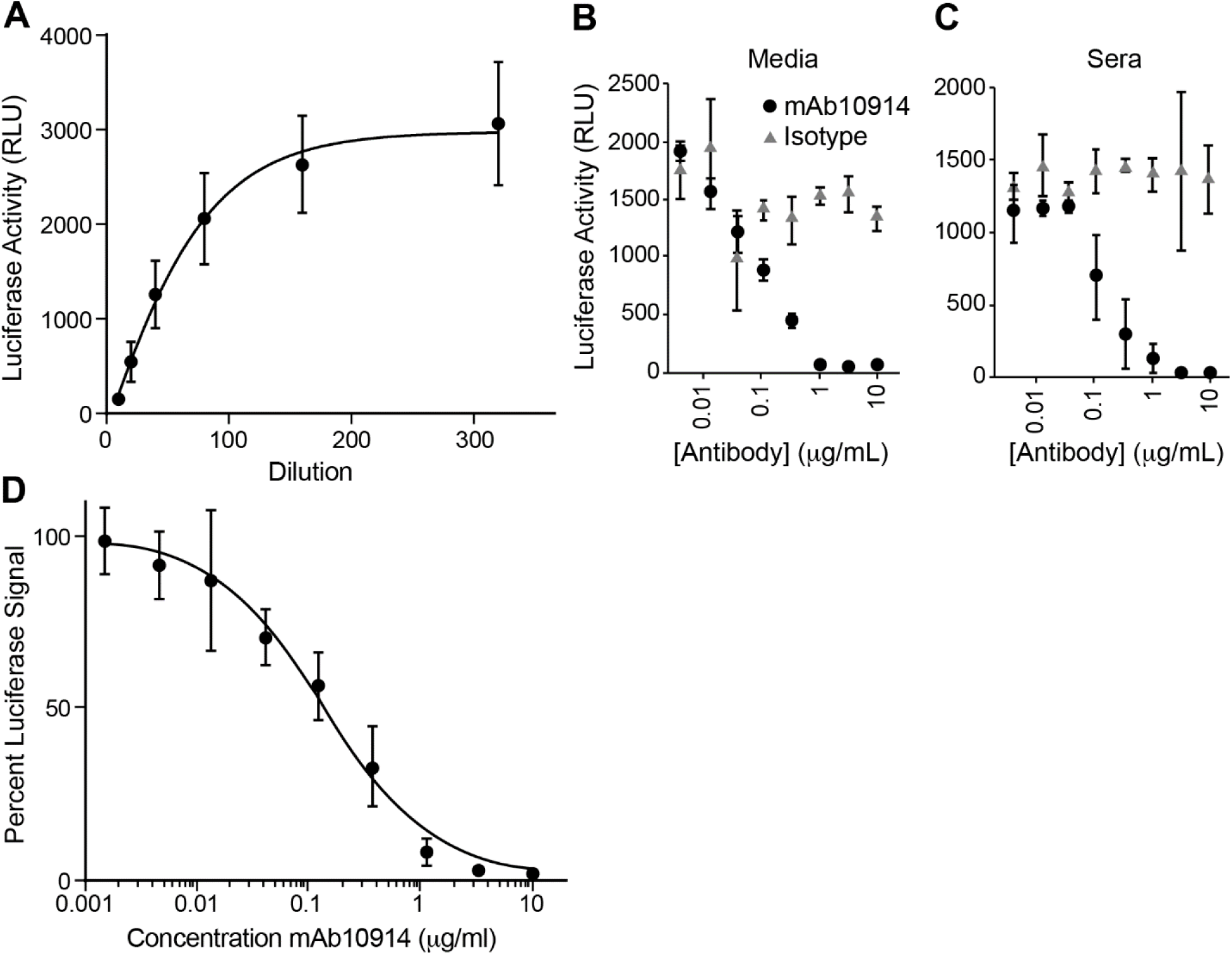
Development of a contrived positive control. A) Minimum recommended dilution. Thirty-nine sera from presumed SARS-CoV-2 seronegative individuals were serially diluted and mixed with VSV-SARS-CoV-2-S-Δ19CT in singlet. Dilutions represent serum dilution in the virus/serum mix. After 30 min at room temperature, the virus/serum mixes were overlaid on Vero-DSP1/Vero-DSP2 monolayers. Luciferase substrate EnduRen™ was added to wells and luciferase activity was measured 22 h after infection. Values represent the average (mean) RLU ± standard deviation, fit with a non-linear regression curve. B&C) Neutralization of VSV-SARS-CoV-2-S-Δ19CT by mAb10914. mAb10914 or isotype control antibody were diluted in OptiMEM (B) or pooled SARS-CoV-2 seronegative sera (C) and incubated with VSV-SARS-CoV-2-S-Δ19CT. Concentrations represent the antibody concentration in the virus/antibody mixes; sera concentration was 1:100. After 30 min at room temperature, the virus mixes were overlaid onto Vero-DSP1/Vero-DSP2 monolayers. EnduRen™ was added to wells and luciferase activity was measured 24 h after infection. Values represent the average (mean) RLU ± standard deviation from duplicate wells. D) Establishment of contrived positive control values. mAb10914 was diluted in pooled SARS-CoV-2 seronegative sera as described in panel C and assayed as in panel C in triplicate runs performed by three different operators. Diluted pooled negative sera was used as a control (0 µg/mL mAb10914). Percent signal relative to the control was determined for each sample. Values represent the average (mean) percent signal ± standard deviation from three independent experiments performed in duplicate, fit with a non-linear regression curve.

### Development of a contrived positive control

Due to the limited availability of large quantities of qualified SARS-CoV-2 seropositive sera, we developed a contrived positive control for the assay. To this end, we spiked various concentrations of mAb10914, a monoclonal antibody that binds to the receptor binding domain of the SARS-CoV-2 spike protein, into pooled SARS-CoV-2 seronegative serum or media alone. As a control, isotype antibody was also used. As expected, mAb10914 caused dose-dependent neutralization of VSV-SARS-CoV-2-S-Δ19CT in both media and pooled negative sera (Figure 6B&C). In contrast, the isotype control antibody did not significantly alter virus-induced luciferase activity, indicating that neutralization by mAb10914 was specific to spike binding. To establish appropriate concentrations to use for contrived controls in validation studies, we tested a total of nine 3-fold serial dilutions of mAb10914 (from 10 µg/mL through ∼0.0015 µg/mL) in duplicate on three separate runs performed by three separate analysts. Because of plate-to-plate variability in raw RLU (relative light unit) values, we normalized results as percent luciferase signal relative to the luciferase signal in control wells containing pooled SARS-CoV-2 seronegative sera. The average (mean) percent luciferase signal at each concentration was plotted, and a non-linear regression curve was generated (Figure 6D). The linear range of the assay was between approximately 10% to 80% luciferase signal or 0.01 to 0.6 µg/mL mAb10914. We selected 2 µg/mL mAb10914 as a contrived positive control high (CPC High), which consistently reduces signal to < 20%. Based on a 95% confidence of the response curve, the concentration of mAb10914 that should elicit a signal below 50% was calculated as 0.18 µg/mL, which thus represents an approximate limit of detection of the assay.

### Examination of assay specificity and sensitivity

Having established a CPC High value, we used it to define assay specificity and sensitivity. To measure specificity, we tested thirty presumptive SARS-CoV-2 seronegative samples alone (matrix) or spiked with 2 µg/mL (CPC High) of mAb10914. This evaluation was conducted to verify that at the optimized 1:100 serum dilution no false positives were obtained and that all contrived positive controls showed virus neutralization. In two separate analytical runs performed by different analysts, no false positives or negatives were observed (Table 1). One negative sample was resulted as indeterminant in the first assay due to one replicate having signal < 50%. However, in the second assay the sample was correctly identified as negative. Thus, in this validation assay specificity was 100%.

**Table 1:** Assay Specificity – Unblinded Analysis

|  | Analyst A |  | Analyst B |  |
| --- | --- | --- | --- | --- |
|  | Diluted Matrix | CPC spike* | Diluted Matrix | CPC spike |
| **Total Tested** | **30** | **30** | **30** | **30** |
| Number Positive | 0 | 30 | 0 | 30 |
| Number Negative | 29 | 0 | 30 | 0 |
| Number Indeterminant | 1† | 30 | 0 | 0 |

\* CPC spike: mAb10914 spiked into diluted matrix sample at 2 µg/mL.
† Sample identified as indeterminant due to one replicate with signal <50%.

To evaluate assay sensitivity, we performed a similar but single-blinded analysis using the same 30 presumptive SARS-CoV-2 seronegative samples. Duplicate samples were assayed, where one replicate sample was spiked with mAb10914 at 2 µg/mL (CPC High) or 0.35 µg/mL (borderline inhibition), before being assigned blinded IDs. Of the 30 un-spiked samples, 26 were identified as negative and 4 as indeterminant (Table 2). We detected 10 of the 11 CPC High samples as positive, and sample preparation error was suspected in the single positive not identified. From the 19 borderline samples, 13 were identified as positive, 4 as negative, and 2 as indeterminant, concordant with the samples containing only a low-level of neutralizing antibody and 0.35 µg/mL being close to the previously estimated assay limit of detection.

**Table 2:** Assay Sensitivity – Blinded Analysis

|  | Sample Type* |  |  |
| --- | --- | --- | --- |
|  | Negative | Borderline | Positive |
| **Number of samples** | **30** | **19** | **11** |
| Result positive | 0 | 13 | 10 |
| Result negative | 26 | 4 | 1† |
| Result indeterminant§ | 4 | 2 | 0 |

\* Sample Type: Negative is diluted seronegative (matrix) sample; borderline is diluted matrix spiked with mAb10914 at 0.35 µg/mL; positive is diluted matrix spiked with mAb10914 at 2 µg/mL.
§ Indeterminant: one replicate with signal >50%, other replicate with signal <50%.
† Technical error during sample preparation is suspected.

### Demonstration of Clinical Agreement

Given that our assay performed well with the contrived controls, we proceeded with blinded studies to determine whether results correlated well with clinical symptoms. To this end, we acquired serum samples from negative donors (symptomless, negative for SARS-CoV-2 by PCR, or SARS-CoV-2 seronegative by ELISA), positive donors (positive for SARS-CoV-2 by PCR or SARS-CoV-2 seropositive by ELISA), and contact positive donors (not tested, but in direct contact with someone positive for SARS-CoV-2 by PCR). A total of 230 samples were received and blinded to the assay analysts until after testing was complete. Of the 125 negative samples received, all 125 tested negative in the assay (Table 3), providing further evidence of our assay’s excellent specificity. All 38 of the known SARS-CoV-2 seropositive samples also tested positive in our assay. Of the 34 samples obtained from donors that had previously tested positive for SARS-CoV-2 by PCR, neutralizing antibodies were identified in 31 samples. Upon closer examination, one sample was collected only 8 days after positive PCR test and was expected to be negative in our assay due to insufficient time for a strong antibody response to develop. From donors who had contact with someone positive for SARS-CoV-2, we identified 13 of 33 samples (39.4%) as positive for neutralizing antibodies. Taken together, our data indicated that our assay has excellent clinical agreement.

**Table 3:**
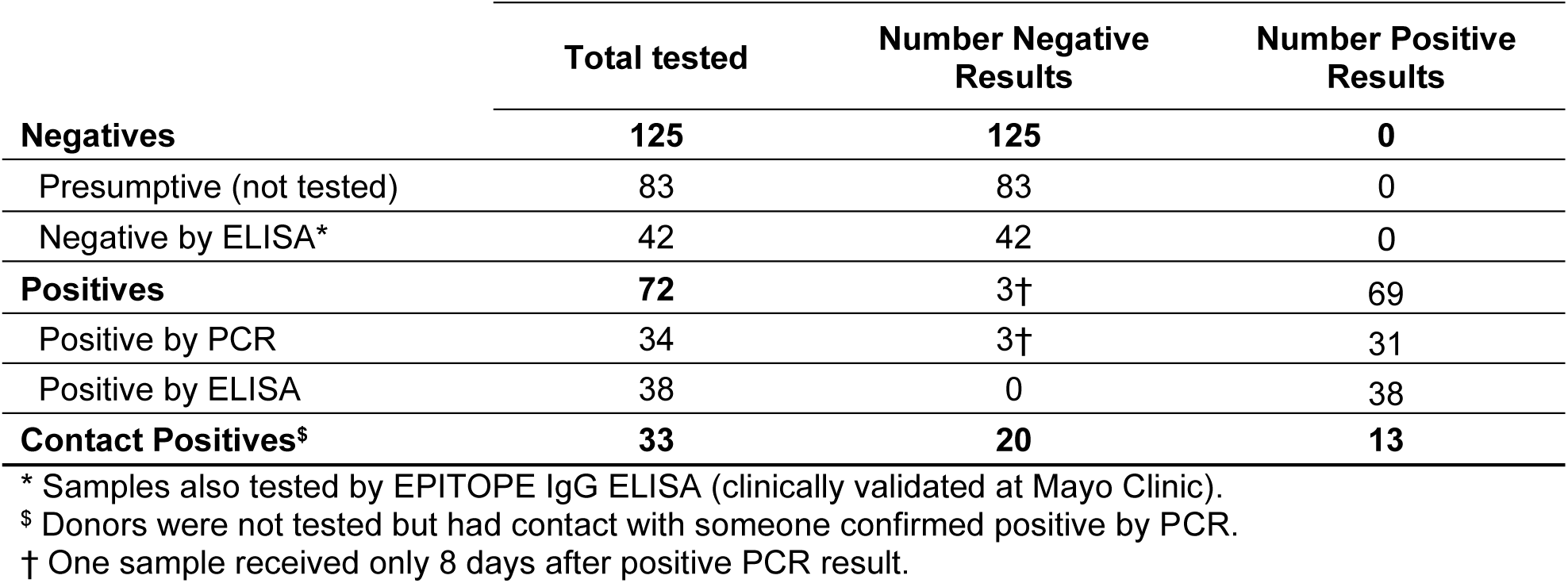
Clinical Correlation – Blinded Studies

### Correlation between VSV-SARS-CoV-2-S-Δ19CT and SARS-CoV-2 neutralization

The strong clinical agreement observed for our assay supported our premise that neutralization of VSV-SARS-CoV-2-S-Δ19CT accurately reflects neutralization of SARS-CoV-2. In order to directly demonstrate this correlation, twenty of the serum samples from above were also tested against a clinical isolate of SARS-CoV-2 (22) in a BSL-3 plaque reduction neutralization test (PRNT). Each of the 5 samples that tested negative in our assay, including the sample acquired eight days after a SARS-CoV-2 positive PCR test, were also below the limit of detection in the PRNT assay (Table 4). Likewise, the 15 samples that tested positive in our assay were also positive in the PRNT assay. Moreover, quantitative virus neutralizing units (VNUs) determined in our assay correlated closely with PRNT_EC50_ values (Table 4 and Figure 7A). To calculate VNUs, a calibration curve consisting of a five-point dilution series of mAb10914 in 1:100 pooled SARS-CoV-2 seronegative sera was included on each assay plate. Based on percent luciferase signal, the equivalent mAb10914 concentration for each sample was determined and multiplied by 100 to give the VNUs for that sample. A strong correlation (Pearson R = 0.8902, p < 0.0001) was observed between VNUs and PRNT_EC50_ values. We also examined the relationship between VNU and virus neutralization titer (VNT) EC50 values. Five 2-fold serial dilutions of forty-four SARS-CoV-2 positive samples were prepared and tested in our assay. The percent luciferase signal for each dilution was plotted and the resulting curves were used to determine the VNT_EC50_ values. Statistical comparison of the VNU and VNT_EC50_ indicated excellent correlation (Spearman R_S_ = 0.8327, p < 0.0001) between the two values (Figure 7B), providing additional demonstration of the accuracy of the VNU values. We concluded that neutralization of VSV-SARS-CoV-2-S-Δ19CT closely replicates neutralization of SARS-CoV-2 and that VNUs provide an accurate method to quantitate the strength of the anti-SARS-CoV-2 antibody response.

**Table 4:** Correlation between IMMUNO-COV and SARS-CoV2 Plaque Reduction Neutralization Test

| Random ID | IMMUNO-COV Result | IMMUNO-COV VNU* | PRNT<sub>EC50</sub>† | Clinical Correlation |
| --- | --- | --- | --- | --- |
| RID-165 | Negative | N/A | < 20<sup>§</sup> | Presumed Negative |
| RID-152 | Negative | N/A | < 20 | Presumed Negative |
| RID-121 | Negative | N/A | < 20 | Contact positive |
| RID-195 | Negative | N/A | < 20 | Contact positive |
| RID-239 | Negative | N/A | < 20 | PCR positive 8-days before testing |
| RID-142 | Positive | 31.8 | 340.0 | PCR Positive |
| RID-168 | Positive | 69.9 | 277.7 | Contact tested positive |
| RID-272 | Positive | 95.2 | 319.8 | PCR Positive |
| RID-288 | Positive | 125.5 | 530.7 | Contact tested positive |
| RID-144 | Positive | 134.2 | 665.8 | PCR Positive |
| RID-138 | Positive | 281.7 | 2061 | Contact tested positive |
| RID-193 | Positive | 345.4 | 1545 | PCR Positive |
| RID-183 | Positive | 402.6 | 1449 | PCR Positive |
| RID-167 | Positive | 430.7 | 845.6 | Contact tested positive |
| RID-287 | Positive | 447.8 | 2130 | PCR Positive |
| RID-290 | Positive | 491.0 | 2999 | Contact tested positive |
| RID-192 | Positive | 775.7 | 4362 | Contact tested positive |
| RID-194 | Positive | 1504.9 | 1502 | Contact tested positive |
| RID-235 | Positive | 2046.6 | 7311 | PCR Positive |
| RID-236 | Positive | 2702.5 | 5203 | PCR Positive |

\* VNU: Virus Neutralization Unit. Arbitrary quantitative value assigned based on calibration curve; higher VNUs represent higher neutralizing antibody levels.
<sup>§</sup> The limit of detection for this assay is 20.
† PRNT<sub>EC50</sub>: Plaque Reduction Neutralization Titer EC50. See Materials and Methods.

**Figure 7:**
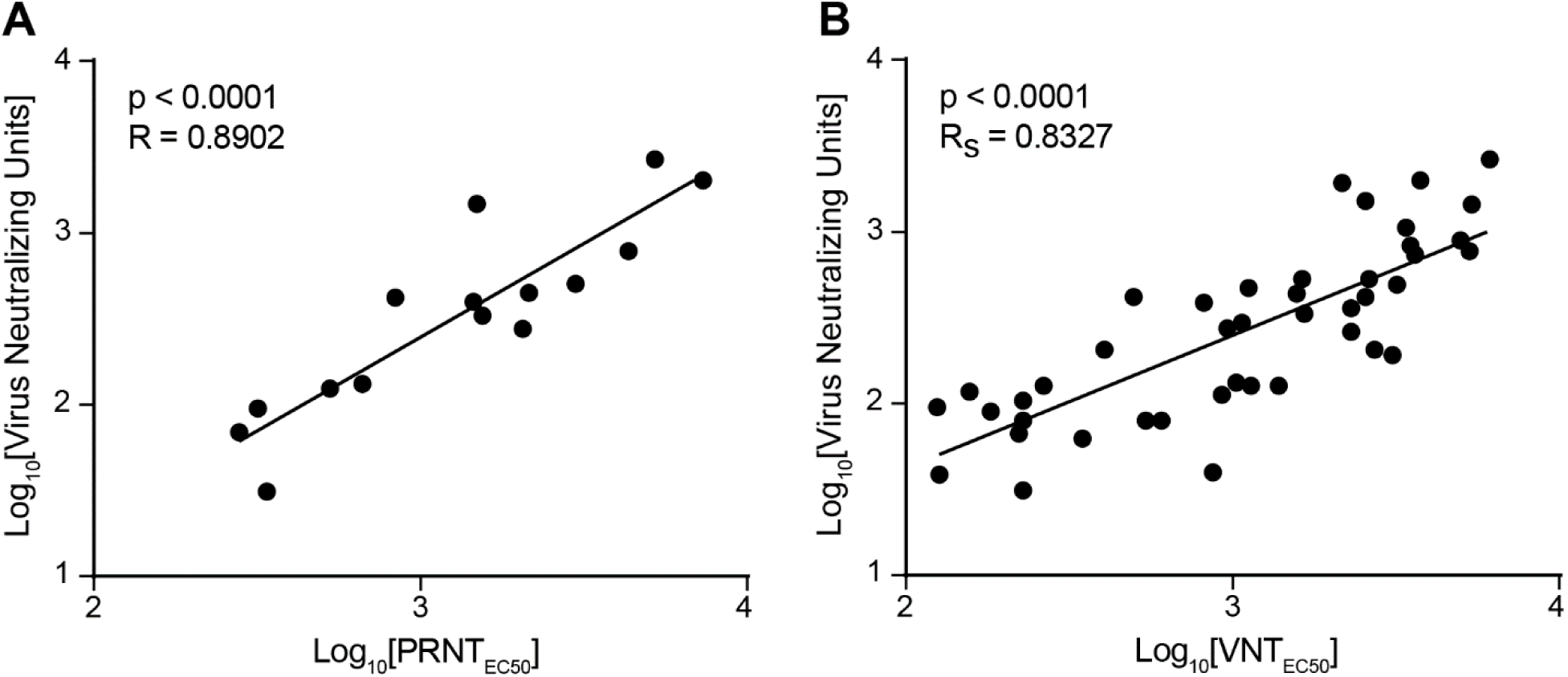
Correlation of virus neutralizing units to PRNT_EC50_ and VNT_EC50_. SARS-CoV-2 seropositive sera samples were serially diluted in OptiMEM and mixed with VSV-SARS-CoV-2-S-Δ19CT. After 30 min at room temperature, the virus/serum mixes were overlaid onto Vero-DSP1/Vero-DSP2 monolayers. Luciferase substrate EnduRen™ was added to wells and luciferase activity was measured between 23 and 27 h after infection. Each assay plate also included a calibration curve in which mAb10914 was spiked into pooled SARS-CoV-2 seronegative sera and mixed with virus. The concentrations of mAb10914 in the virus mixes for the calibration curve were 3, 1, 0.33, 0.11, and 0.037 µg/mL. The percent signal for each test sample and calibration curve point were determined. A virus neutralizing unit (VNU) was determined for each sample based on its percent signal relative to the calibration curve, where a VNU equals the concentration of mAb10914 for the given percent signal multiplied by 100. A) Comparison with PRNT_EC50_. Of the positive samples tested, 15 were subjected to a plaque reduction neutralization test (PRNT) using a clinical isolate of SARS-CoV-2. Two-fold serial dilutions of samples were tested in the PRNT50 assay from 1:20 through 1:40960. For each sample, the number of plaques at each dilution was plotted and used to determine the PRNT EC50 value for the sample. Statistical comparison of VNU relative to PRNT_EC50_ was performed. Data was transformed due to non-normal distribution (p < 0.0001). B) Comparison with VNT_EC50_. The percent signal for each sample dilution was plotted and used to determine the VNT_EC50_. Statistical comparison of VNU relative to VNT_EC50_ was performed. Data was transformed due to non-normal distribution (p < 0.0001).

## Discussion

Numerous ELISA and rapid diagnostic tests have been developed to detect anti-SARS-CoV-2 antibodies, and several are approved in various countries for diagnostic testing. Yet, an easily scalable clinical assay for detecting antibodies capable of neutralizing SARS-CoV-2 has remained relatively elusive. In particular, studies of vaccine efficacy and evaluations of convalescing plasma therapies require high-throughput quantitative testing for anti-SARS-CoV-2-neutralizing antibodies. Here, we describe our assay IMMUNO-COV™ for high-throughput, quantitative, and clinically-validated measurement of SARS-CoV-2-neutralizing antibodies at low biocontainment (BSL2). We predict IMMUNO-COV™ will play a key role in screening donor samples for SARS-CoV-2 convalescing plasma therapy trials and evaluating immune responses to candidate vaccines, as well as offering further insights into SARS-CoV-2 immunity.

Our assay exploits the fusion phenotype of recombinant VSV-SARS-CoV-2-S-Δ19CT to produce quantitative luciferase signal in 96-well plate format, where virus neutralization by test serum samples is detected as a reduction in luciferase activity (Figure 1). Importantly, results from IMMUNO-COV™ correlated closely with plaque reduction neutralization EC50 values (p < 0.0001) determined using a clinical isolate of SARS-CoV-2 (Table 4 and Figure 7A). Thus, neutralization of VSV-SARS-CoV-2-S-Δ19CT accurately reflects neutralization of SARS-CoV-2. Recently, strong correlation was also observed between neutralization of other SARS-CoV-2 spike-expressing VSVs and SARS-CoV-2 (12,13), providing further evidence that inhibition of replicating VSVs encoding SARS-CoV-2 spike mimics SARS-CoV-2 neutralization.

IMMUNO-COV™ is a validated laboratory developed test. In validation and verification studies, the assay had 100% specificity, and no false positives were detected in any of the testing performed (Tables 1, 2, and 3). All samples that tested positive by ELISA also tested positive by IMMUNO-COV™, which provided additional quantitation of the neutralizing antibody levels. Two samples collected from donors that previously tested PCR positive for SARS-CoV-2 were negative in our assay. Because medical histories were self-reported, we were unable to verify the accuracy of the PCR test results. Yet regardless of previous SARS-CoV-2 status, negative results in the IMMUNO-COV™ assay indicate that the neutralizing antibody response is weak. Both samples were collected less than 3 weeks from the self-reported SARS-CoV-2 positive test, and certain factors can dampen or delay the development of robust antibody responses. Moreover, neutralizing antibody responses represent a gradient of strengths. Because it is not known at which point on the continuum an anti-SARS-CoV-2 antibody response becomes adequately neutralizing to be considered protective, it is important to have quantitative measures of neutralizing antibody responses, and it is difficult to conclusively define an assay sensitivity. Using a borderline contrived control, 68.4% of samples were positive for SARS-CoV-2-neutralizing antibodies (Table 2). However, based on agreement with clinical symptoms, assay sensitivity was at least 97.2% in tested serum samples (Table 3).

Additional validation studies are ongoing to define the long-term stability of critical reagents and matrix samples. Early results from these studies are promising and indicate that serum samples are stable under a variety of storage conditions and are not affected by up to three freeze-thaw cycles. While our formal validation did not include plasma samples, experiments with NL1 (Figure 4B, Figure 5B, and data not shown) indicated IMMUNO-COV™ is also compatible with plasma samples.

Traditionally, assays that measure virus neutralizing antibodies rely on a series of serial sample dilutions to quantitate the level of anti-viral antibodies present in each test sample as a virus neutralizing titer or EC50 neutralizing titer (VNT_EC50_). A unique feature of IMMUNO-COV™ is that quantitation of virus neutralization is performed without a full sample dilution series. By testing samples at fewer dilutions, assay throughput is increased, and up to 41 samples can be assayed in duplicate on a single 96-well plate with necessary controls (see example plate layout in Figure 8). To improve assay sensitivity, testing is performed at the minimum recommended dilution, which we determined during validation studies to be 1:100 (Figure 6). At this dilution, approximately 83% of the samples tested fell within the linear range, though approximately 38% of positive samples fell above the linear range. To extend the quantitation range for samples containing high levels of SARS-CoV-2 neutralizing antibodies, a second dilution, perhaps 1:1000, can be tested in a follow-up assay.

**Figure 8:**
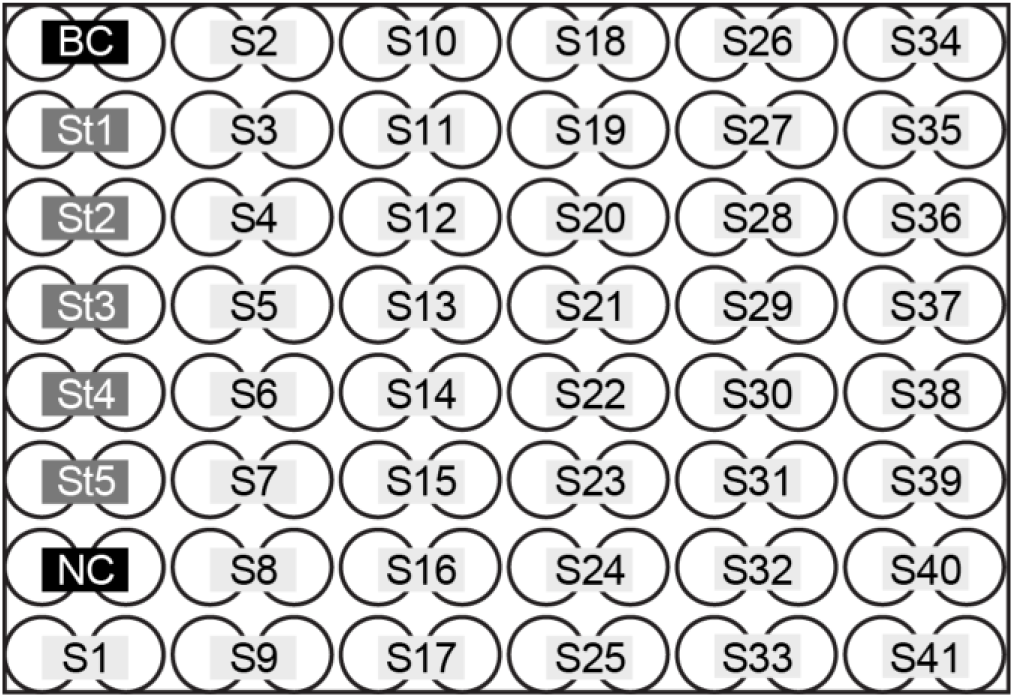
Example assay plate layout. All controls and samples are assayed in duplicate. A total of 41 samples (S1 to S41) can be run on a single plate using a single dilution for each sample. BC (background control) consists of pooled SARS-CoV-2 seronegative serum at 1:100 without virus. NC (negative control), consists of pooled SARS-CoV-2 seronegative serum at 1:100 with VSV-SARS-CoV-2-S-Δ19CT. NC is used to determine 100% luciferase signal in the assay. St1 through St5 are standards for the calibration curve. All standards are pooled SARS-CoV-2 seronegative serum at 1:100 with VSV-SARS-CoV-2-S-Δ19CT and mAb10914 spiked into each standard at different concentrations: Std1 is 3 µg/mL, Std2 is 1 µg/mL, Std3 is 0.33 µg/mL, Std4 is 0.11 µg/mL, and Std5 is 0.037 µg/mL.

Virus neutralization units (VNUs) are the unit of quantitation used in the IMMUNO-COV™ assay. VNUs are calculated as 100X the amount of mAb10914 that yields the same percent luciferase signal as the test sample based on a standard curve. The quantifiable range for seropositive VNUs is approximately 20 to 200, which corresponds to concentrations of 0.2 to 2 µg/mL mAb10914. Because VNUs are an arbitrary value, it is useful to compare them to VNTs when interpreting results in the context of traditional assays. VNUs and VNT_EC50_ values correlated well (p < 0.0001) in the IMMUNO-COV™ assay (Figure 7B), providing a means for relating VNUs to VNTs. As additional test samples are analyzed and data generated, the comparison of VNUs and VNTs will be further refined.

Quantification of the strength of neutralizing antibody responses is absolutely required to accurately assess immune responses to new SARS-CoV-2 vaccines. Strong antibody responses are required for protective immunity, but concerns have also arisen regarding potential antibody-dependent enhancement of disease. Studies with MERS and SARS-CoV have demonstrated antibody-dependent enhancement of infection *in vitro* (16,23–25). Studies in non-human primates also suggest possible disease augmentation with increasing antibody levels for SARS-CoV (26), and some evidence supports antibody-dependent enhancement in clinical cases of SARS (27,28). Yet early studies using COVID-19 convalescing plasma to treat severe cases of COVID-19 have shown great promise (3–5). How the ratio of total anti-SARS-CoV-2 antibodies to SARS-CoV-2 neutralizing antibodies may shift potential outcomes between disease enhancement and resolution is not fully understood. Nevertheless, it is widely agreed that strong neutralizing antibody responses are desirable for both vaccine responses and convalescent plasma therapies. IMMUNO-COV™ offers the capacity for widespread clinical testing of neutralizing antibody levels, which can accelerate evaluation and further development of candidate vaccines and convalescing plasma therapies. Thus, IMMUNO-COV™ is a valuable addition to the currently available repertoire of clinical tests.

## Materials and Methods

### Cells

African green monkey Vero cells (ATCC® CCL-81™), Vero-αHis (29), and baby hamster kidney BHK-21 cells (ATCC® CCL-10™) were maintained in high-glucose DMEM supplemented with 5% fetal bovine serum and 1X penicillin/streptomycin (complete media) at 37°C/5% CO_2_. To generate Vero-DSP1-Puro (Vero-DSP1) and Vero-DSP2-Puro (Vero-DSP2) were generated by transducing Vero cells with lentiviral vectors SFFV-DSP_1-7_-P2A-Puro or SFFV-DSP_8-11_-P2A-Puro encoding the Renilla Rluc8 luciferase-GFP split reporter DSP1 or DSP2 (GenBank EF446136.1) (20) under control of the spleen focus forming virus (SFFV) promoter and linked to the puromycin resistance gene via a P2A cleavage peptide. Transduced cells were selected using 10 µg/mL puromycin. Following selection, Vero-DSP cells were maintained in complete media supplemented with 5 µg/mL puromycin. Puromycin was excluded when cells were seeded for assays.

### Generation of VSV-SARS-CoV-2-S-Δ19CT

Full-length human codon-optimized SARS-CoV-2 Spike (S) glycoprotein (NC_045512.2) in pUC57 was obtained from GenScript (MC_0101081). The plasmid was used as a PCR template to generate a cDNA encoding SARS-CoV-2 Spike with a deletion in the nucleotides encoding the C-terminal 19 amino acids (S-Δ19CT) and 5’ MluI and 3’ NheI restriction sites. To generate the viral genome, the amplified PCR product containing S-Δ19CT was cloned into pVSV-M51R in place of VSV-G using the MluI and NheI restriction sites (Figure 2A). Plasmid was sequence verified and used for infectious virus rescue on BHK-21 cells as previously described (30). VSV-G was co-transfected into the BHK-21 cells to facilitate rescue but was not present in subsequent passages of the virus. The virus was propagated in Vero-αHis cells by inoculating 80% confluent monolayers in 10-cm plates with 1 mL of virus. Viruses were harvested 48 hours after inoculation, aliquoted, and stored at −80°C until use. Aliquots were used for tissue culture infectious dose 50 (TCID_50_) assay on Vero-αHis cells.

### Infections

Vero monolayers in 6-well plates were inoculated with OptiMEM alone (mock) or 2 mL of passage 3 VSV-SARS-CoV-2-S-Δ19CT in OptiMEM. After 4 h at 37°C/5% CO_2_, fresh OptiMEM was added to wells. Cells were returned to the 37°C/5% CO_2_ incubator until 20 h after infection, when they were photographed at 100x magnification using an inverted microscope.

### Reagents

EnduRen™ live cell substrate was prepared according to the manufacturer’s instructions (Promega #E6482). For inhibition studies, mouse α-SARS-CoV-2 spike antibody (GeneTex #GTX632604), affinity-purified polyclonal α-human-Ace-2 antibody (R&D Systems #AF933), and recombinant human Ace-2 (R&D Systems #933-ZN-010) were diluted in OptiMEM to the desired concentrations. mAb10914 is a human α-SARS-CoV-2 Spike neutralizing monoclonal antibody that was prepared and scaled up using methods previously described (31) by Regeneron Pharmaceuticals, Inc. (manuscript submitted).

### Luciferase Assays

Assays were performed in 96-well black-walled plates with clear bottoms. EnduRen™ substrate was added to wells (final concentration of 7.5 µM) at various times after infection, but at least 2 hours before reading bioluminescence on a Tecan Infinity or Tecan M Plex instrument (2000 ms integration, 200 ms settle time). For kinetic experiments, repeated bioluminescence reads were performed without the addition of more EnduRen™ for up to 24 hours after initial EnduRen™ addition; at later time points, additional EnduRen™ was added to wells at least 2 hours prior to reading bioluminescence.

### Collection of plasma and sera samples

A clinical protocol to collect blood samples for assay validation was reviewed and approved by Western IRB (study ID: VYR-COV-001). Serum samples were collected from patients who had previously tested positive for SARS-CoV-2 infection by a PCR test, from patients who had known exposure to individuals infected with SARS-CoV-2 and symptoms of COVID-19, as well as a cohort of patients of were presumed negative for antibodies, i.e. individuals who never had symptoms of COVID-19 disease and were likely never exposed. Clinical information was self-reported. A total of 150 adults were enrolled at BioTrial based in Newark, New Jersey and Olmsted Medical Center in Rochester, Minnesota. Eighty additional sera aliquots from samples that had been tested using the EPITOPE IgG ELISA (validated assay) were obtained from Mayo Clinic, Rochester MN.

### Blinded sample testing

Sera samples were assigned a random ID prior to being given to analysts for testing. Samples were assayed in batches, with an unknown number of positive and negative samples in each batch. Each sample was assayed at a 1:100 dilution, with all positive and some negative samples being further assayed in a dilution series including 1:100, 1:200, 1:400, 1:800, 1:1600, and for some samples 1:3200.

### PRNT Assay

Sera was heat-inactivated for 30 min at 56°C and serially diluted 2-fold in Dulbecco’s minimal essential medium supplemented with 2% heat-inactivated fetal bovine serum. SARS-CoV-2 (USA-WA1/2020) (22) was diluted to approximately 200 PFU/mL and mixed with an equal volume of diluted sera (final dilutions of serum with virus were 1:20, 1:40, 1:80, 1:160, 1:320, 1:640, 1:1280, 1:2560, 1:5120, 1:10240, 1:20480, 1:40960). Virus mixed with an equal volume of media alone was used as a control. After a 1 h incubation at 37°C, 250 µL of virus/serum or virus/media mixes were used to inoculate Vero-E6 monolayers in 6-well plates. Inoculations proceeded for 1 h at 37°C with occasional rocking, before monolayers were overlaid with 4 mL of 1.6% low-melting agarose in Minimal Essential Media supplemented with 4% fetal bovine serum and antibiotics. Plates were incubated at 37°C for 2 days when plaques appeared, then fixed with 10% formaldehyde, and stained with 2 mL of 0.05% neutral red, incubated for 6 h at 37°C. Plaques were counted and the number at each dilution was plotted and used to determine the PRNT EC50 value for each sample by a nonlinear regression model. Plaque counts greater than 30 were too numerous to count and were considered as equivalent to the virus/media control.

### Flow cytometry

Vero-DSP1-Puro cells were dislodged using Versene, counted, and transferred to microcentrifuge tubes (5×10^5^ cells/tube was used for Ace-2 staining and 1.5×10^6^ cells/tube was used for TMPRSS2 staining). For Ace-2 staining, cells were pelleted and resuspended in 100 µL FACS buffer (2% FBS in DPBS) containing 0.2 µg goat-α-human Ace-2 (R&D Systems #AF933). After 30 min on ice, cells were rinsed with 1 mL FACS buffer and resuspended in 100 µL FACS buffer containing 5 µL donkey-α-goat IgG-PE secondary antibody. After 30 min on ice, cells were rinsed with 1 mL FACS buffer and fixed with 1% paraformaldehyde for 15 min on ice. Cells were washed twice with FACS buffer, resuspended in 500 µL FACS buffer and analyzed on a CYTOFLEX flow cytometer (Beckman Coulter). For TMPRSS2 staining cells were resuspended in 1 mL ice-cold 70% ethanol in DPBS and incubated on ice for 10 min. Cells were centrifuged, washed once with 1 mL FACS buffer, and resuspended in 100 µL of a 0.5% saponin solution containing 4 µg rabbit α-TMPRSS2 (Invitrogen #PA5-14264). After 30 min on ice, samples were washed twice with 1 mL FACS buffer and resuspended in 100 µL of a 0.5% saponin solution containing 2 µL goat α-rabbit IgG-AF647 secondary antibody. After 30 min on ice, cells were washed twice with FACS buffer and fixed with 1% paraformaldehyde for 15 min on ice. Cells were washed twice with FACS buffer, resuspended in 500 µL FACS buffer and analyzed on a CYTOFLEX flow cytometer (Beckman Coulter). For both Ace-2 and TMPRSS2 staining, secondary antibody only controls were used for isotype.

### Immunoblot

Total cells and culture supernatants of Vero cells infected with VSV-GFP or VSV-SARS-CoV-2-S-Δ19CT or mock-infected were harvested 48 h after infection. Virus was released from the cell membranes by freeze-thaw and cell debris was removed by centrifugation. Samples were diluted in duplicate in LDS sample buffer (Invitrogen #B0007) and reducing agent (Invitrogen #B0009) according to the manufacturer’s directions, incubated at 70°C for 10 min, and 40 µL of each sample was ran on duplicate 4-12% Bis-Tris gels (Invitrogen #NW04125Box) along with precision plus protein dual color standard (Bio-Rad #161-0374). Proteins were transferred to nitrocellulose membranes using a Power Blotter XL. Membranes were blocked in 5% non-fat dry milk in TBST, washed three times with TBST, and incubated for 1 h at room temperature with primary antibody mouse α-SARS-CoV-2 Spike (1:1000, GeneTex #GTX632604) or rabbit polyclonal α-VSV (1:5000, Imanis Life Sciences #REA005). Membranes were washed three times with TBST and incubated for 1 h at room temperature with secondary antibody goat α-mouse IgG-HRP (Prometheus #20-304) or goat α-rabbit IgG-HRP (Prometheus #20-303) at 1:20,000. Membranes were washed three times with TBST, and protein bands were developed for 2 min at room temperature using ProSignal® Dura ECL Reagent (Prometheus #20-301). Protein bands were imaged using a BioRad ChemiDoc Imaging System.

### Determination of percent luciferase signal and virus neutralization units

Raw relative light unit (RLU) values for each well were blank corrected using the mean RLU value from background wells receiving pooled SARS-CoV-2 seronegative serum at 1:100 but no virus. The percent signal was determined for each sample well as the corrected RLU value divided by the mean corrected RLU value from negative control wells receiving virus and pooled SARS-CoV-2 seronegative serum at 1:100. Samples with a percent signal < 50% for both sample replicates were classified as positive. Samples with a percent signal ≥ 50% for both sample replicates were classified as negative. When one replicate exhibited a positive response and the other a negative response, the sample was classified as indeterminant and testing was repeated. Virus neutralizing titers (VNUs) were determined for positive samples based on a calibration curve. The calibration curve was run on each plate and consisted of mAb10914 spiked into pooled SARS-CoV-2 seronegative sera at 3, 1, 0.33, 0.11, and 0.037 µg/mL. From the calibration curve, the equivalent concentration of neutralizing antibody for a given percent signal was determined. To convert to VNU, the antibody equivalent concentration was multiplied by 100.

### Statistical Analyses

Descriptive statistics, comparisons, and regression analyses were performed in Graph Pad Prism, v8.4.0 (San Diego, CA). Tests for normality of variance were conducted, and whenever possible parametric comparisons were used. For non-normal datasets, non-parametric approaches were used. A four-parameter non-linear regression was used for the calibration curve of the virus neutralizing units within the assay. For correlation analyses, Spearman’s or Pearson’s correlation analysis was conducted.

## Acknowledgements

We would like to thank all the participants in our clinical trial; without their willingness to donate blood we would not have been able to develop and validate this assay. We would also like to thank the entire Imanis Life Sciences, Vyriad, and Regeneron teams for helpful discussions, laboratory, and operational support. Natalie Thornburg provided the SARS-CoV-2 isolates and Kenneth and Jessica Plate provided virus stocks, cells, and protocols. Funding for this project was provided by Regeneron Pharmaceuticals, Inc. as part of an ongoing collaboration with Vyriad, Inc., and by NIH grant AI120942 to SCW.

